# Health Conditions Associated with Overweight in Climacteric Women

**DOI:** 10.1101/661827

**Authors:** Maria Suzana Marques, Ronilson Ferreira Freitas, Daniela Araújo Veloso Popoff, Fernanda Piana Santos Lima de Oliveira, Maria Helena Rodrigues Moreira, Andreia Maria Araújo Drummond, Dorothéa Schmidt França, Luís Antônio Nogueira dos Santos, Marcelo Eustáquio de Siqueira e Rocha, João Pedro Brant Rocha, Maria Clara Brant Rocha, Maria Fernanda Santos Figueiredo Brito, Antônio Prates Caldeira, Fabiana Aparecida Maria Borborema, Viviane Maria Santos, Josiane Santos Brant Rocha

## Abstract

This study aims to investigate the association between health conditions and overweight in climacteric women assisted by primary care professionals. It is a cross-sectional study conducted with 874 women from 40 to 65 years of age, selected by probabilistic sampling between August 2014 and August 2015. In addition to the outcome variable, other variables such as overweight/obesity, sociodemographic, reproductive, clinical, eating and behavioural factors were evaluated. Descriptive analyses of the variables investigated through their frequency distributions were performed. Then, bivariate analyses were performed through Poisson regression. For the multiple analyses, the hierarchical Poisson regression was used to identify factors associated with overweight/obesity in the climacteric period. The prevalence of overweight/obesity was 74%. Attending public school (PR: 1.30 - 95% CI 1.14 - 1.50), low schooling (PR: 1.11 - 95% CI 1.01 - 1.23), gout (PR: 1.18 - 95% CI 1.16-1.44), kidney disease (PR: 1.18 - 95% CI 1.05 - 1.32), metabolic syndrome - MS (PR: 1.19 - 95% CI 1.05 - 1.34) and fat intake (PR: 1.12 - 95% CI 1.02 - 1.23) were considered risk factors for overweight. Having the first birth after 18 years (PR: 0.89 - 95% CI 0.82 to 0.97) was shown to be a protective factor for overweight and obesity. The presence of overweight and obesity is associated with socio-demographic, reproductive, clinical and eating habits.

## Introduction

Brazil has been presenting a rapid process of demographic and epidemiological transition, leading to the frequent occurrence of chronic-degenerative diseases^1^. The increase in the prevalence of overweight, represented by overweight and obesity, among the elderly female population raises great concern in developed and developing countries. Since overweight and obesity are risk factors for adverse health events^2^ such as disturbances in lipid and glycidic metabolism, psychological stress and sleep alterations, with increasing risk of cardiovascular diseases^3^, musculoskeletal disease, acute myocardial infarction^4^, cancer^5^ and worse quality of life in comparison to those who were satisfied with their body weight ^6^.

Overweight and obesity have become a public health problem in the world. The projection for 2025 is that about 2.3 billion adults are overweight, and more than 700 million are obese. According to a study conducted in 2016, the rate of overweight among Brazilian women is 50.5%, increasing this frequency with age and up to 64 years^7^.

Epidemiological data are still scarce associating excess weight with behavioural and clinical variables in climacteric women^8^, using probabilistic samples^9^. Considering that climacteric is an important period of the women life cycle, and that this period is related to the potential peak of fat mass and obesity in this group, the current study aimed to investigate the association between health conditions and excess of weight ratio in climacteric women, assisted by primary care professionals, since this phase may assume a pathological character or be associated with other chronic diseases.

## Materials and methods

A cross-sectional and analytical study was carried out in the city of Montes Claros, Minas Gerais, Brazil, from August 2014 to August 2015, whose target population consisted of 30,801 climacteric women enrolled in 73 health care units, excluding pregnant, postpartum or bedridden women. This study was carried in the Family Health Strategy (FHS) that represents the mechanism of primary health care (PHC) on the public health system in Brazil ^10^.

Sampling was of the probabilistic type, and the selection of the sample occurred in two stages. Each health care unit team was taken as a conglomerate, being drawn 20 units, covering the urban and rural area for data collection. Following this, a proportional number of women were randomly selected according to the climacteric stratification criteria of the Brazilian Society of Climacteric (SOBRAC), in 2013^11^. Women between 40 and 65 years of age, enrolled in the selected teams and physically able to respond to the questionnaires and submitted to anthropometric measurements and laboratory tests (12-hour fasting) were considered eligible to participate in the study. The researchers made the previous training of all collectors and interviewers and maintained supervision during the data collection stage. After this selection, the women were invited to present themselves for the research in a previously established date. The final sample consisted of 874 climacteric women who were invited to sign the Informed and Post-Informed Consent.

The nutritional status of women was evaluated by body mass index (BMI), which was considered as a study outcome. Despite the inclusion of some patients over 60 years, women were categorised into eutrophic (BMI <25 kg / m^2^) and overweight (IMC ≥ 25 kg/m^2^), following a categorisation model used in other studies with similar population groups ^12, 13, 14^. Initially, women were weighed wearing light clothing and without footwear, in orthostatic position, with their feet together and arms relaxed throughout the body, by a mechanical anthropometric medical scale (Balmak 11) with a capacity of 150 kg and divided into 100 g. The stature was measured by anthropometer (SECA 206^®^), fixed in a flat wall and without skirting. In this measurement, the women were instructed to keep their feet together, in an upright position, with their head positioned in the Frankfurt plane. For the calculation of BMI, the body weight in kilograms was divided by the squared height, expressed in meters (BMI = P/A^2^).

The women answered questions related to the independent variables, which were allocated in three blocks: (1) sociodemographic, (2) reproductive (3) clinical, eating and behavioural habits.

The block of sociodemographic variables included age (40-45, 46-51, 52-65 years); type of school (public or private); level of schooling (elementary school I, elementary school II, high school / higher education); marital status (married, separated, divorced, widowed); labor occupation (yes or no); monthly income (≥ 01 minimum wage, <01 minimum wage), being the minimum wage equivalent to US$217,42 at the time of data collection; number of people residing in the same house (up to 2, more than 2) and; skin color (white, not white).

The reproductive variables comprised the age of menarche (≤ 11 years, 12-14 years and ≥ 15 years), first birth weight (<4000g; ≥ 4000g), climacteric symptoms assessed by the Kupperman Index^15^ (absent/mild; moderate/severe) and age of first delivery (≤ 18 years old> 18 years).

The clinical, eating and behavioral variables included: liver disease (absent, present), gout (absent; present), renal disease (absent; present), Metabolic Syndrome (MS) (absent; present); Urinary incontinence (absent, present), cardiovascular disease risk (low risk, intermediate risk, high risk), drinking (yes, no), fat intake (yes, no), smoking (yes or no), symptoms of depression, quality of sleep and physical activity.

Metabolic Syndrome (MS) was evaluated using the NCEP-ATPIII criteria of the Brazilian Society of Diagnosis and Treatment of MS^6^, urinary incontinence was assessed by the International Consultation on Incontinence Questionnaire-Short Form ICIQ-SF^17^, the risk for cardiovascular diseases was assessed by Framingham Global Risk Score^18^, the symptoms of depression were evaluated by the Beck Depression Inventory^19^, sleep quality was assessed by the Pittsburgh Sleep Quality Index^20^ and physical activity practice was assessed through the International Physical Activity Questionaire (IPAQ short version) ^21^.

The women were submitted to peripheral venous blood collection to analyse the laboratory parameters. Serum triglyceride levels were determined by the colourimetric enzymatic method. The level of HDL (High-Density Lipoprotein) cholesterol was obtained by selective precipitation of LDL (Low-Density Lipoprotein) cholesterol and VLDL (Very Low-Density Lipoprotein) cholesterol with dextran sulphate in the presence of magnesium ions, followed by dosing by enzymatic system cholesterol oxidase/peroxidase with calorimetry and reading, as performed in the total cholesterol dosage, using Labtest^®^ reagents, in Cobas Mira^®22^ apparatus. The lipid profile was analysed according to parameters proposed by the Brazilian Society of Cardiology^23^ and fasting glycemia according to the standards of the Expert Committee on the Diagnosis and Classification of Diabetes Mellitus^24^.

The data was tabulated in the statistical software Statistical Package for Social Science (SPSS, version 21, Chicago, Illinois). Initially, descriptive analyses of all variables through their frequency distributions were carried out, and then, bivariate analyses of the outcome-variable with each independent variable were performed using the Chi-Square Test. Gross Prevalence Ratios (PR) were estimated with their respective 95% confidence intervals. Variables with a descriptive level (p-value) of less than 0.25 were selected for multiple analysis using the hierarchical Poisson regression model, adapted to the model proposed by other authors^9^. The model was composed of blocks of variables at distal (sociodemographic variables), intermediate (reproductive) and proximal (clinical, eating and behavioural) variables. Prevalence ratios (PR) adjusted with their respective 95% confidence intervals were estimated, remaining in the model only those that presented descriptive level p<0.05. At each hierarchical level, the stepwise forward procedure was adopted: starting the model with the statistically significant variable selected in the bivariate analysis, and then adding other variables (Fig. 1).

**Figure 1:**
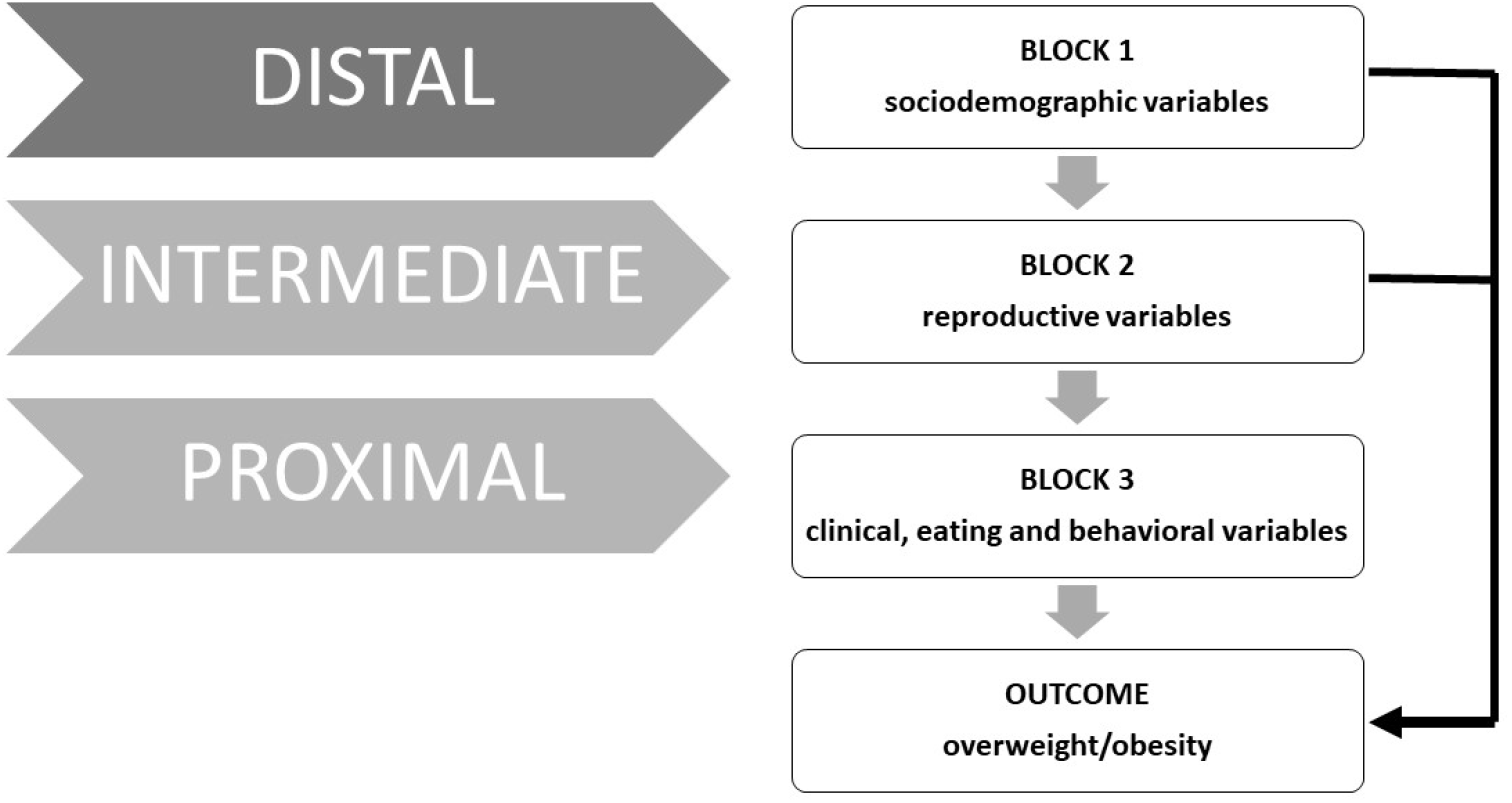
Model with the statistically significant variable selected in the bivariate analysis, and then adding other variables.

As this study involved humans, it was submitted, appreciated and approved for execution by an Ethics and Research Committee (No. 817,166), with the ethical precepts of the Brazilian Resolution CNS 466/2012 being fully observed.

## Results

The sample consisted of 874 women between 40 and 65 years of age, of whom 74.1% were overweight. When categorised by climacteric phases, it was observed that postmenopausal women had a higher prevalence of overweight/obesity (54.3%).

The results of the bivariate analysis revealed the following variables associated with the overweight/obesity outcome: age between 52 and 65 years (p = 0.184), private school (p = 0.000), low schooling (p = 0.093) (p = 0.0006), liver disease (p = 0.000), gout (p = 0.000), disease (p = 0.106), weight of the 1st child at birth equal to or greater than 4000 g (p = .039), high risk for cardiovascular diseases (p = 0.000), alcohol consumption (p = 0.039) and fat intake (p = 0.065). However, women between 46 and 51 years of age (p = 0.184), who had a late menarche age (p = 0.039) and had children over 18 years old (p = 0.004) had a protective effect against overweight and obesity. It should be emphasized that there was a high prevalence of overweight and obesity in all the independent variables presented (Table 1).

**Table 1:**
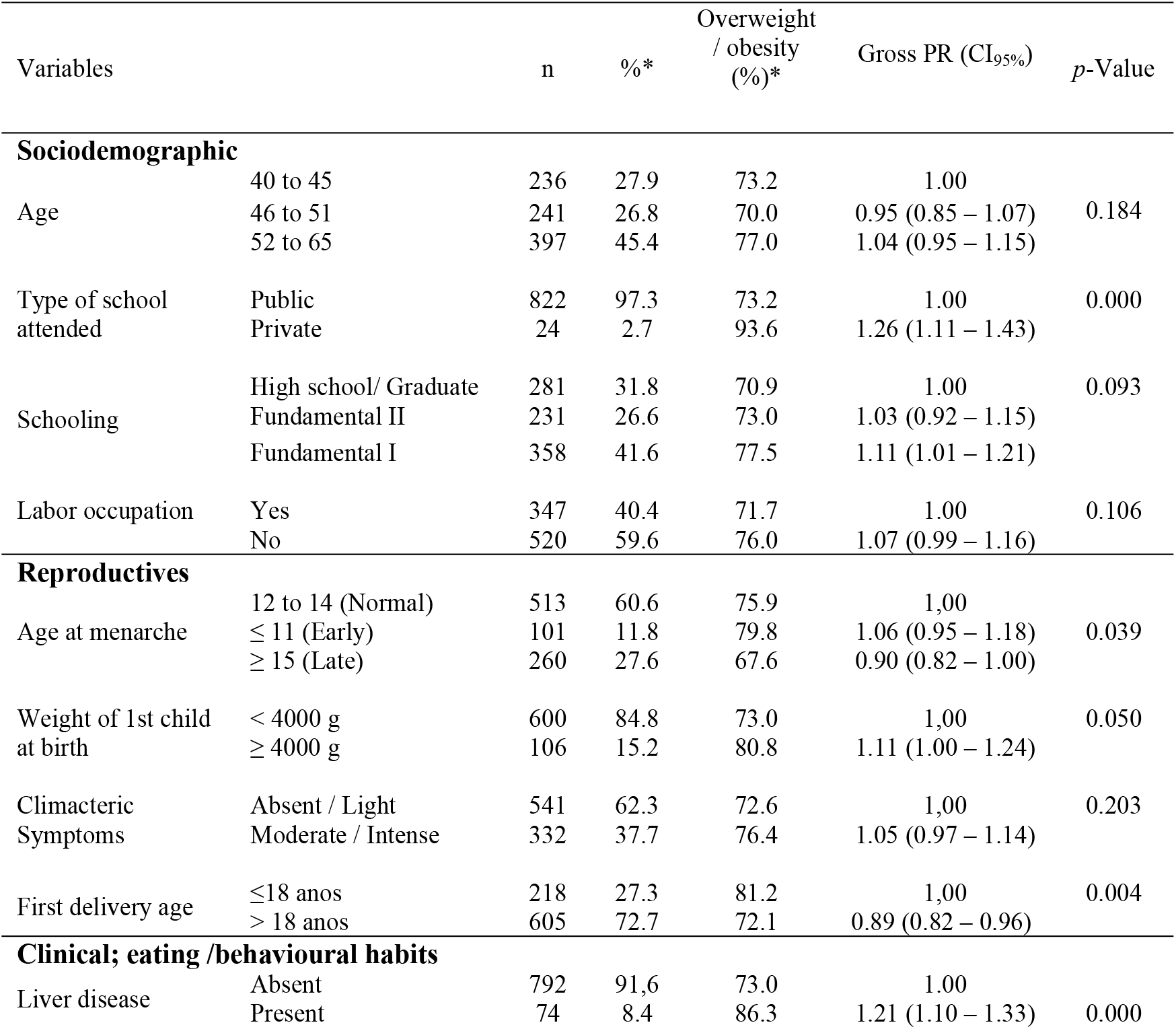
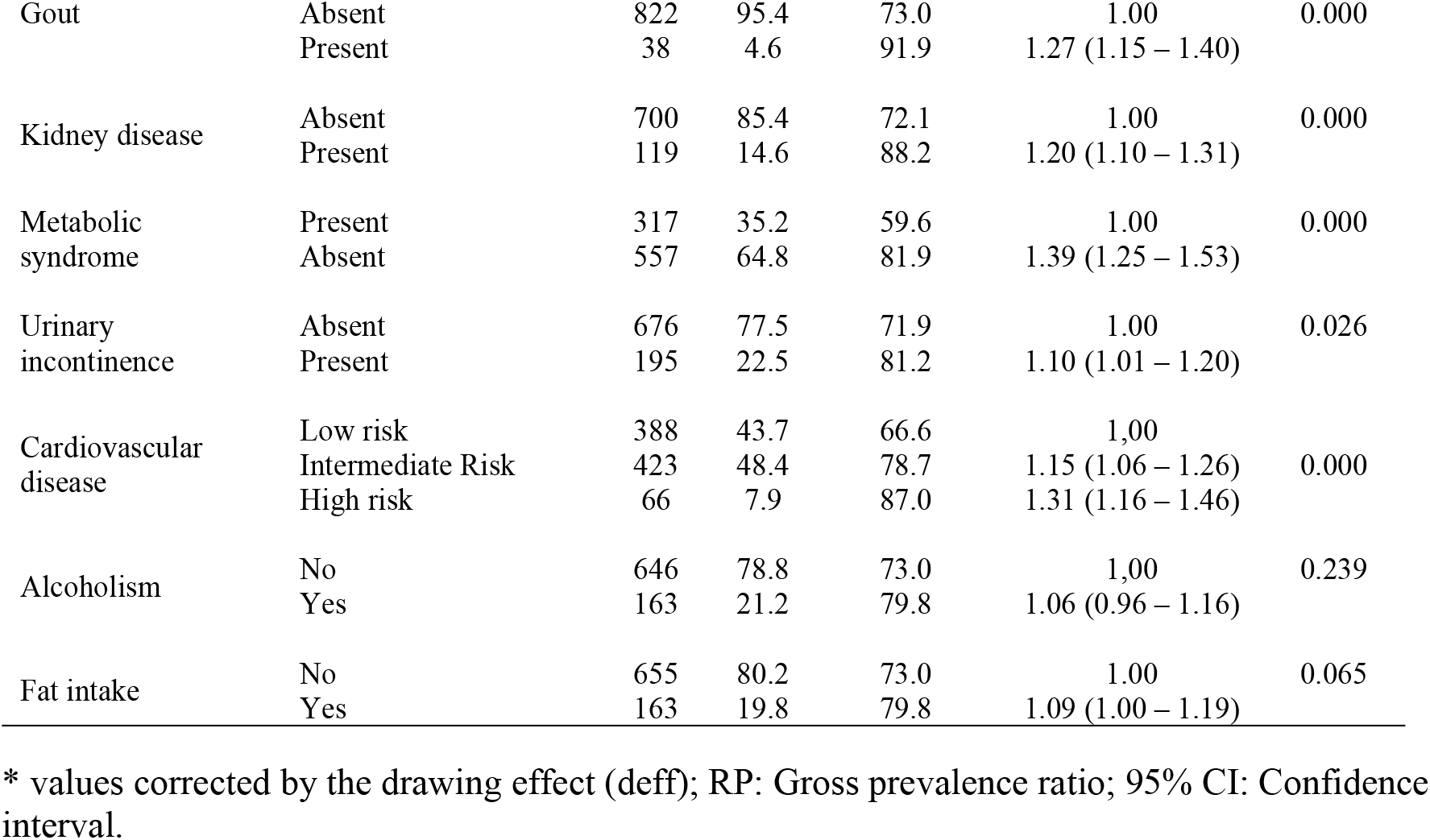
Sample characteristics and gross prevalence ratio (PR) for overweight/obese according to sociodemographic factors, reproductive, clinical, behavioural and eating habits of menopausal women.

The socio-demographic (marital status, monthly income, number of individuals residing in the same house and color of skin), clinical and behavioral (smoking, physical activity, depression symptoms, sleep quality) factors did not present significant associations (p <0.250) with overweight/obesity, and are not included in the hierarchical model.

The health conditions that were associated with overweight/obesity in the hierarchical model at the distal level were private school (PR = 1.30, p = 0.000) and low level of education (PR = 1.11, p = 0.033). After adjusting for sociodemographic factors, an association at intermediate level between the first childbirth above 18 years (PR = 0.90, p = 0.010) was observed, and this variable had a protective effect against the occurrence of overweight/obesity (Table 2). At the proximal level, after adjusting for the potential confounding factors analyzed, the presence of gout (RP = 1.18, p = 0.004), MS (RP = 1.29, p = 0.000), kidney disease (P = 1.19, p = 0.006) and fat intake (PR = 1.12, p = 0.014) were found to be positively associated with overweight/obesity (Table 2).

**Table 2:** Prevalence ratio adjusted for overweight/obesity according to sociodemographic, reproductive, clinical, eating and behavioural factors of climacteric women.

| Variables |  | PR (CI<sub>95%</sub>)<br>adjusted | *p* Value |
| --- | --- | --- | --- |
| **Sociodemographic (distal level)** |  |  |  |
| Type of school attended | Public | 1.00 |  |
|  | Private | 1.30 (1.14 -1.50) | 0.000 |
| Schooling | High School/Graduate | 1.00 |  |
|  | Fundamental II | 1.05 (0.94 – 1.17) | 0.420 |
|  | Fundamental I | 1.11 (1.01 -1.23) | 0.033 |
| **Reproductives (Intermediate level)** |  |  |  |
| First delivery age | ≤18 anos | 1.00 |  |
|  | > 18 anos | 0.90 (0.82 -0.97) | 0.010 |
| **Clinical; eating/behavioural and habits (proximal level)** |  |  |  |
| Gout | Absent | 1.00 |  |
|  | Present | 1.18 (1.05 -1.32) | 0.004 |
| Metabolic Syndrome | Absent | 1.00 |  |
|  | Present | 1.29 (1.16 -1.44) | 0.000 |
| Kidney disease | Absent | 1.00 |  |
|  | Present | 1.18 (1.08 -1.29) | 0.000 |
| Cardiovascular disease | Low risk | 1.00 |  |
|  | Intermediate risk | 1.05 (0.95–1.15) | 0.332 |
|  | High risk | 1.19 (1.05–1.34) | 0.006 |
| Fat intake | No | 1.00 |  |
|  | Yes | 1.12 (1.02 -1.23) | 0.014 |

PR: Gross and adjusted prevalence ratio; 95% CI: confidence interval

## Discussion

The prevalence of overweight and obesity in the population of the present study was higher than 2/3 of the sample, with a BMI mean of 28.67 ± 6.35 kg/m^2^, with a predominance of overweight in postmenopausal women. These findings are in accordance with a study conducted in São Paulo/Brazil, where the BMI mean in postmenopausal women was 29.0 ± 5.6 kg/m^2^ ^25^.

Weight gain in climacteric is due to the ageing process and estrogenic depletion, with the centralised distribution of fat mass related to ovarian failure ^26^, which leads to a change in the hormonal environment previously dominated by estrogen to an environment where there is a predominance of testosterone, favouring androgenicity ^27^. Also, inadequate lifestyle habits, such as sedentary lifestyle, consumption of fats and sugars, can lead to physiological and metabolic alterations^28^. The limited perception of body weight and the importance of the control^29^, and the use of medications such as antidepressants, analgesics, and anxiolytics ^30^ also compete for the appearance of this condition.

Obesity is associated with insulin resistance and chronic inflammation predisposing to various diseases, including breast cancer, whose pathogenesis has been linked to increased estrogen levels^31^.

In addition, excessive body weight also contributes to the occurrence of systemic arterial hypertension (SAH), depression and worsening of climacteric symptoms^32^. Together with other comorbidities, they impair the quality of life of women and impact their functionality^33^.

According to the findings of the study, having attended private school seems to be associated with overweight in the climacteric. This may be due to the more accessible food with high caloric rates in childhood and adolescence, leading to weight excess, which could be perpetuating in adult life. However, the literature cannot explain these findings consistently, presenting evidence of a higher prevalence of weight excess among students of private schools in other age groups ^34;35^.

Still, some studies evidence the association between low schooling and high BMI^36^, in consonance with the present findings, suggesting that a higher level of education may favour healthier living habits, such as the intake of vegetables and fruits^37^ and the regular practice of physical activity^38^.

Regarding the gynaecological aspects, having a delivery occurred after the age of 18 was shown to be a protective factor for overweight/obesity. Other studies have also shown an association between overweight/obesity, early parturition and parity^39,40^. Findings suggest that younger maternal age at first delivery is independently associated with a higher risk of central obesity and MS in climacteric^41^. One explanation would be the possibility of a higher number of pregnancies among women with early parturition and lifestyle changes, although the pathophysiology of this association is still unclear and deserves additional studies^42^. Multiparity is associated with an increase in the prevalence of MS since it favours abdominal obesity^43^ and insulin resistance in climacteric women^44^.

The diagnosis of gout and overweight/obesity in climacteric are also associated. This finding becomes relevant since hyperuricemia correlated with insulin resistance, hypertension, obstructive sleep apnea, chronic renal disease (CKD), MS and elevated cardiovascular risk^45,46^. According to this context, hyperuricemia may be related to an increase in the prevalence of coronary artery disease (CAD) and to the incidence of major cardiovascular events in climacteric women, being an independent risk factor^47^. Chromosomal abnormalities are associated with elevated serum levels of uric acid and gout in postmenopausal women, demonstrating a possible role of sex hormones in the regulation of the urate transporter in gout^48^.

An association between kidney disease and overweight was found in the present study. These data are consistent with the Brazilian Society of Nephrology’s Dialysis Survey in 2014, which showed that 37% of dialysis patients were overweight or obese as a risk factor for CKD^49^. In addition, obesity is associated with MS, that is also a risk factor for the development of CKD^50^.

Overweight is related to compensatory hyperfiltration, which occurs to meet the metabolic demands increased by body weight, with possible damage to the kidneys and increased risk of long-term glomerulopathy, besides being also a risk factor for nephrolithiasis and kidney cancer. The obese patient also has a higher relative risk for developing albuminuria and a decrease in the glomerular filtration rate, even without CKD^51^.

In climacteric, with increased risk for obesity, MS becomes more prevalent, increasing the incidence of cardiovascular disease and the risk of acute myocardial infarction (AMI)^52^, a vulnerability attributed to the decrease of estrogen and insulin resistance^53^. The association between overweight/obesity and MS was observed in the present study with the consequent risk elevation for cardiovascular diseases. Another study corroborated these findings and demonstrated that the prevalence of MS was also higher in postmenopausal women^54^. Obesity presents as a possible primary factor for the occurrence of MS and the risk of cardiovascular diseases, since an overweight patient may also have visceral adiposity, which is one of the diagnostic criteria of MS.

Among the overweight/obese women in this study, the diet characterised by fat intake was associated with overweight. A document published by the Health Surveillance Agency points out that excessive consumption of saturated fat, as well as sugars, is related to the development of chronic noncommunicable diseases, including obesity^55^. The balance in fat consumption is a viable strategy for a possible reduction of cardiovascular risk in this population^56^, since inadequate diet is the leading cause of cardiovascular mortality^23^.

The present study presents as limiting factors the use of BMI as the sole diagnostic criterion for overweight/obesity, in detriment to the use of other gold standard techniques, such as dual X-ray densitometry (DEXA). The variables liver diseases, kidney disease and gout, were addressed by self-report, and it was not possible to establish with precision the different etiologies of these diseases, but being able to establish their association in a generic way, provoking the need for further studies using more accurate diagnostic tools, such as imaging or laboratory tests. Moreover, it is a cross-sectional study and, therefore, unable to establish causality among the studied variables. Despite the presented limitations, the obtained results bring relevant information on the subject, besides listing variables to be studied in future researches. It should be emphasised that the sample used in the study was representative of the population and was obtained in a probabilistic way, strengthening the results and associations obtained.

In addition, from a socioeconomic point of view, the population studied resides in a region that represents the Brazilian reality with confidence, located in a transition zone between considered rich Brazil (represented by the South and Southeast states) and Brazil with characteristics of poverty (represented by the Northern and Northeastern states). Therefore, the present study brings relevant associations on the health of climacteric women in an emblematic and representative area of the Brazilian population.

## Conclusion

The presence of overweight/obesity was associated with climacteric women who had attended private schools, with low schooling, gout, metabolic syndrome, kidney disease, with high cardiovascular risk and who ingest fats in the diet. In turn, having first delivery after 18 years of age was presented as a protective factor for women not to become overweight/obese. It is suggested to monitor the modifiable factors since they were associated with overweight in climacteric women assisted by primary health care services.

